# Identifying stem cell numbers and functional heterogeneities during post-embryonic organ growth

**DOI:** 10.1101/2020.04.08.032110

**Authors:** Diana-Patricia Danciu, Julian Stolper, Lázaro Centanin, Anna Marciniak-Czochra

## Abstract

Uncovering the number of stem cells necessary to grow an organ has been challenging in most vertebrate systems. Here, we have developed a mathematical model that we use to characterise stem cells in the fish gill, an organ that displays non-exhaustive growth. Our work employs a Markov model, first stochastically simulated via an adapted Gillespie algorithm, and further improved by using probability theory. The stochastic algorithm produces a simulated data set for comparison with experimental data by inspecting quantifiable properties, while the analytical approach skips the step of *in silico* data generation and goes directly to the quantification, being more abstract and very efficient. By applying the model to a large clonal experimental dataset, we report that a reduced number of stem cells are responsible for growing and maintaining the fish gill. The model also highlights a functional heterogeneity among the stem cells involved, where activation and quiescence phases determine their relative growth contribution. Overall, our work presents an easy-to-apply algorithm to infer the number of stem cells functionally required in a life-long growing system.

## Introduction

Stem cells are essential during organ growth in all higher vertebrates. Once the organisms reach maturation and acquire a definitive body size, their adult stem cells (aSCs) are responsible for maintaining homeostasis, i.e. they continue to proliferate with the goal of replacing the cells lost regularly. Unlike mammals and other higher vertebrates, fish increase their body size throughout their entire life. Accordingly, fish organs must adapt to this permanent growth, by either increasing in size as happens with gills (Stolper et al., 2019) and retina (Centanin et al., 2014; Tsingos et al., 2019), or in numbers as is the case with neuromasts - mechano-sensory organs sensing the water flow (Ghysen and Dambly-Chaudière, 2007; Dambly-Chaudière et al., 2003; Wada et al., 2013; Seleit et al., 2017). Fish aSCs carry the task of organ remodelling, and they are not only able to maintain homeostasis, but also to drive growth. In this work, we focus on the respiratory organ of fish - the gill or branchia, recently introduced as a suitable model for studying aSCs and organ development (Stolper et al., 2019). The model organism under study is the Japanese rice fish (*Oryzias latipes*), colloquially known as medaka, which is convenient due to its rapid development (Fig. 1B) and isogenic genome (Wittbrodt et al., 2002). Two different stem cell populations have been reported by Stolper et al. (2019) to drive growth and maintain homeostasis, with the growth stem cells being restricted to the growing edge of the organs.

**Figure 1:**
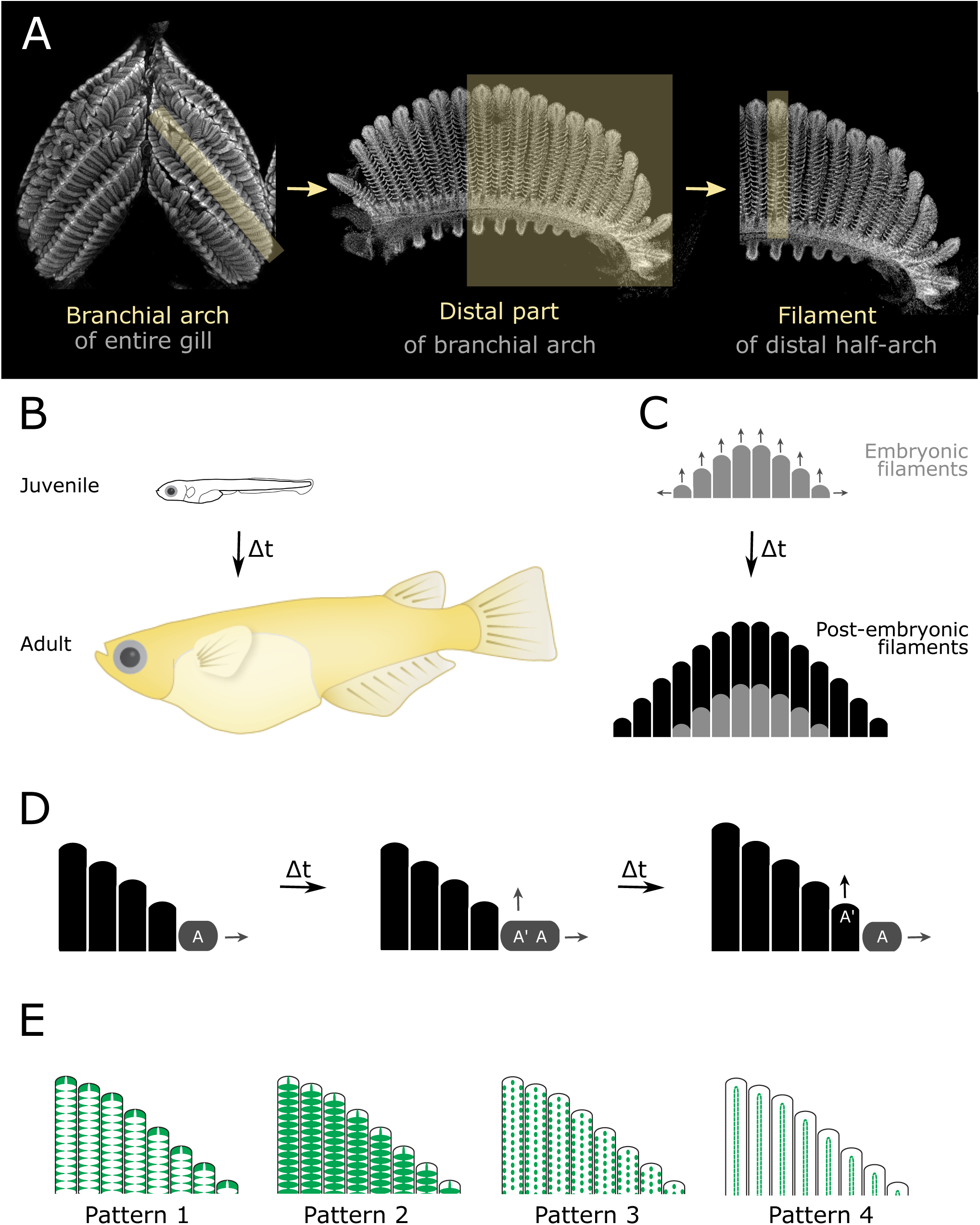
Biological background of medaka gills: **A**. Structure of a gill: entire gill (left) contains 4 pairs of branchial arches (middle), each of which is composed of a sequence of filaments (right). Our study focuses on half-arches (distal and proximal) **B**. Development of medaka fish. **C**. The gills grow along two axes, by elongating filaments and by adding new ones. New filaments are generated by stem cells residing in niches at the basal extremities of branchial arches, while their elongation is driven by stem cells at the tip of each filament. **D**. Simplified scheme for the post-embryonic growth of branchial arches: when a stem cell A in the niche divides (left), its progeny A’ generates a new filament (middle) and it drives its elongation from the tip (right). This process continues throughout the fish life. **E**. Four different cell lineages have been reported (Stolper et al., 2019) which, when labelled, give rise to four different filamental patterns.

Each gill is composed of four pairs of branchial arches, which in turn comprise a sequence of filaments, each of which is built from multiple stacked lamellae (see Fig. 1A and study by Leguen (2018)). Branchial arches grow along two orthogonal axes: longitudinally by elongating their filaments, and transversally by adding new ones (Fig. 1C). Both these actions are performed by stem cells located at the growing edges of the respective domains: growth stem cells at the periphery of branchial arches (*br-archSCs*) generate new filaments, which contain growth stem cells at the tip (*filamSCs*), driving their elongation by adding new lamellae. Homeostatic stem cells are present along the mid-axis of filaments, at the base of each lamella, and are responsible for replenishing the cell population supporting the tissue. Development of branchial arches presents a hierarchical setting: A *br-archSC* divides so that its daughter cell creates the new filament, where it becomes a *filamSC* that drives its elongation from the tip by leaving a trail of homeostatic stem cells along the way, which have the role to maintain the lamellae (Fig. 1D).

Stolper et al. (2019) have recently shown that the growth *br-archSCs* include four different fate restricted stem cells which are recruited together into a newly forming filament, giving rise to four different patterns (Fig. 1E). This means that the different stem cell types coordinate their activity and division to generate a new filament. Such a concept opens many questions and avenues to explore: How do the stem cells coordinate their behaviour in order to work as an ensemble? How do the fate restricted cells get recruited to the new filament and regulate their division so that they maintain the ratio of different cell types within the filament? A first step in approaching such questions is to determine the number of *br-archSCs* of each fate, which are at the base of the hierarchy, being essential for setting in motion the organ development mechanism.

Accordingly, this work develops mathematical tools for counting the stem cells choosing each fate, responsible for generating filaments. Defining the number of stem cells involved in a life-long process has proven difficult in most systems, mainly due to the lack of specific markers. Stem cell counting has been possible in intestinal crypts by the existence of a specific marker (Snippert et al., 2010) and it has been inferred in the hematopoiesis system by using genetic tricks that allow a combinatorial label (Pei et al., 2017; Busch et al., 2015; Sun et al., 2014). In medaka fish gills, stem cell specific markers are not available, so lineage tracing experiments are performed with a ubiquitous labelling, and data are only available at one time point since fish need to be sacrificed. The essential property of fish gills, which facilitates our analysis despite data restrictions, are their modular structure. Therefore the experimental images resulting from lineage tracing experiments can provide a history of *br-archSCs* divisions.

## Results

### Stochastic model suggests that stem cells are not homogeneous in their division behaviour

In our previous paper (Stolper et al., 2019), we have studied the nature of the filament-generating cells, by investigating two different hypotheses to determine their self-renewal capacity. Are these stem cells (SCs), capable of creating multiple filaments, or are these cells a group of progenitors, each of which is capable of producing exactly one filament? We tested each of the two hypotheses, based on clonal data acquired using lineage tracing tools (Gaudi toolkit described by Centanin et al. (2014)) and recording the labelling status of each filament along a branchial arch. The experimental procedure labels a small number of cells at embryonic stages with a nuclear-tagged Green Fluorescent Protein (GFP). Briefly, the Gaudi^*RSG*^ medaka line expresses a Red Fluorescent Protein (RFP) ubiquitously, and this RFP prevents the expression of a nuclear GFP (nGFP). Upon induction with Tamoxifen or after a heat shock, the RFP is removed from the genome allowing nGFP expression. Since this constitutes a modification in the genome of the cell, the same modification will be found in all its progeny, which will be recognised by the nGFP expression.

In the gill system, if a *br-archSC* is labelled with nGFP, then the filaments originating from it will be green as well. And along the same lines, filaments coming from a non-labelled cell will not be green. Accordingly, each branchial arch is described by an array of binary values corresponding to labelled (1) and unlabelled (0) filaments (Fig. 2A and Supplementary Material). Since the focus is on post-embryonic filaments, for each branchial arch, we selected the 8 most peripheral filaments from each half of the arch (from now on referred to as “mini-arches”) and rearranged them so that in the resulting table the first column consists of the oldest filament of the mini-arch, i.e. the 8th filament counted from the *br-archSCs* niche (Fig. 2A). The choice of selecting only 8 filaments is made based on the fact that only branchial arches with more than 25 filaments are considered, and taking into account that normally there exist 5-8 embryonic filaments in the middle of the arch, which were generated before induction of genetic recombination. As previously presented by Stolper et al. (2019), there are at least 4 different fate-restricted *br-archSCs* in the peripheral niche (Fig. 1E). These give rise to different patterns of nGFP that are easily distinguishable in the filaments and were named (Patterns 1-4). Therefore, our data set consists of arrays containing values in the set {0, 1, 2, 3, 4}, or combinations of patterns recorded by values of the form 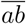 with *a, b* ∈ {1, 2, 3, 4}. Each pattern is analysed separately, so the original data are remodelled into four data sets, one for each pattern, with binary values (Fig. 2A).

**Figure 2:**
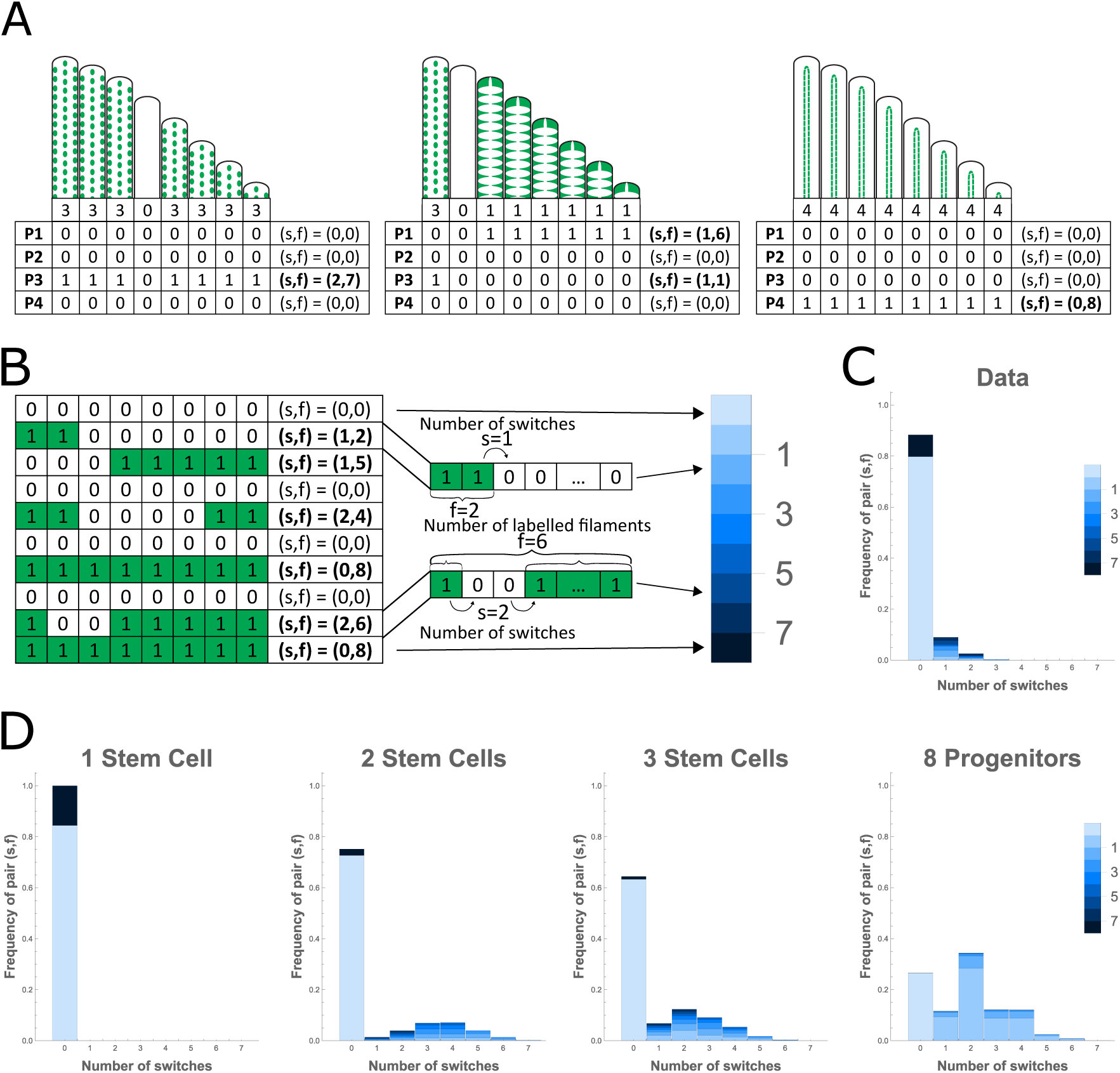
Approach and results of the homogeneous model: **A**. Toy mini-arches used to describe how the data is prepared for the analysis. For each mini-arch, the distribution of patterns is recorded as 1-4 (first row of each table), while mixed patterns are recorded as 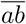, *a, b* ∈ {1, 2, 3, 4} (supplementary table S1). For each pattern, the configuration is rewritten such that each mini-arch is described by an array of binary entries (bottom four rows of the tables, showing the distribution for each pattern P1-P4). Finally, for each such array, the number of switches *s* (i.e. 1-to-0 and 0-to-1 transitions) and that of labelled filaments *f* is determined, and recorded as a pair (*s, f*). **B**. Example of 10 mini-arches extracted from the experimental data, adjusted for studying Pattern 1. For each mini-arch the pair (*s, f*) of switches and labelled filaments is recorded and shown in the last column. Focusing on two mini-arches, the switches are indicated and the labelled filaments are counted, the number of which corresponds to a blue tone on the colour bar shown, which will be used in the following plots. **C**. Plot of the 8-filament long mini-arches in the entire dataset, for Pattern 1. The plot shows the frequency of observing a certain pair (*s, f*) of switches and labelled filaments (y-axis), according to the number of switches (*s*) observed on the x-axis, and the number of labelled filaments (*f*) represented by the colour code. One can notice that most mini-arches are composed of filaments carrying the same label/pattern (0 switches), and out of these, most are fully unlabelled. This ratio between entirely labelled and entirely unlabelled mini-arches is regulated by the experimental labelling efficiency, which is approximated from the data. For pattern 1, the labelling efficiency is 15.53% (i.e. cells have a probability 0.1553 of being labelled). **D**. Comparison between various simulated scenarios. The two extreme scenarios described by Stolper et al. (2019), the stem cell model (left) and the progenitor model (right), are plotted as the data in C, for comparison. The stem cell model, even though extreme, provides a much better fit to the data from C, than the progenitor model does. The central plots show results of simulations for scenarios in which 2 (middle-left) and 3 (middle-right) stem cells reside in the niche, for Pattern 1. Here a homogeneous division behaviour is assumed, i.e. stem cells are randomly chosen for division and filament generation, with all having the same probability of being selected. In such a homogeneous setting, the more stem cells are introduced into the system, the worse the fit to the data becomes, and the progression can be followed from left to right.

For the comparison between models and data, the notion of switches was introduced, defined as the transition from a labelled to an unlabelled filament, i.e. binary transitions. Accordingly, for each mini-arch we counted the switches and the labelled filaments, and recorded them as a pair of the form (*s, f*) with *s* ∈ {0, 1, …, 7} switches and *f* ∈ {0, 1, …, 8} labelled filaments (Fig. 2A-B). For a mini-arch of 8 filaments, out of the total number 8 × 9 of associations (*s, f*) only 33 are possible (for example, for *s* = 0 the only options are *f* ∈ {0, 8}, while for *s* = 7 one must have *f* = 4). The frequency of observing each such pair (*s, f*) in the data can be computed (Fig. 2C).

Stolper et al. (2019) have shown that a stem cell model considering all filaments in the mini-arch to be generated by one single stem cell provides a better fit to the data than a progenitor model in which each filament is created by a different progenitor cell (Fig. 2D, far left and far right, re-considers the two scenarios, for 8-filament long mini-arches, for Pattern 1). But since we often observe switches in the branchial arches, it follows that more than one stem cell are contributing filaments to the branchial arch. Interestingly, the amount of labelled and unlabelled filaments is not identical, nor does it follow a certain distribution, indicating that if more than one stem cell contribute filaments, they are not homogeneous in their probability of division. This insight is supported by the two modelled cases where two and three functionally homogeneous stem cells contribute filaments to the branchial arch, and compared each scenario to experimental data obtained from clonal analysis (middle plots in Fig. 2D). These simulations are performed by first selecting the number of labelled stem cells, *L* ∼ Binomial(*n, probLab*), where *n* is the total number of stem cells in the niche (in the two cases considered in Fig. 2D middle, *n* = 2 or *n* = 3) and *probLab* is the experimental labelling efficiency approximated from the data (e.g. *probLab* = 0.1553 for Pattern 1) via a combinatorial approach (see STAR Methods). The labelling efficiency is different for each pattern and represents an average over the whole data set, which only considers the oldest filament in each mini-arch, as it provides an indication of the first cell which divided post-embryonically. Next, a weighted random choice selects whether a labelled or an unlabelled stem cell will divide, at each time step, with weights given by the number of labelled and unlabelled *br-archSCs*, for each of the eight filaments in the arch. Following this study, we concluded that the filament-generating cells are indeed stem cells (*br-archSCs*), but just having a small number of functionally homogeneous *br-archSCs* contributing filaments to the branchial arches is not sufficient for explaining the data.

### A Markov approach shows that fish gills originate from a small number of functionally heterogeneous stem cells

As opposed to a homogeneous scenario in which *br-archSCs* (both labelled and unlabelled) divide randomly giving rise to an uncorrelated succession of labels, the stretches of consecutive identically labelled filaments observed in the data and the previously tested cases (of two and three homogeneous SCs - Fig. 2D, middle) suggest successive generations (divisions) of the same *br-archSC*. This novel heterogeneity idea is derived and quantified from the data via the mathematical study, as a purely experimental approach could not indicate such a behaviour. The heterogeneous scenario means that not all *br-archSCs* in the niche have equal probabilities of being selected for division at a specific time step, and corresponds to the concept of stem cell activation and quiescence phases, in which when a stem cell divides, it becomes activated and divides multiple times before another SC takes over. The heterogeneity is incorporated in the model through the “probability of division” parameter *p*, representing the probability that the stem cell which has just divided will be the next one to divide again. If this probability *p* = 1, then all filaments in the mini-arch will carry the label of the first stem cell which divided, while a probability of division *p* = 0.5 corresponds to the most homogeneous scenario with intermingled labelled and unlabelled filaments with comparable incidence. This correlation between the cell which has just divided and the one about to divide, for *p* > 0.5, will be referred to as a “heterogeneous division behavior” or a “functional heterogeneity”.

Accordingly, because of the assumption of single-step memory of the system that takes into account which cell divided last, the heterogeneity scenario can be modelled by a Markov process on the generated filaments (Fig. 3A-B). If a labelled filament was generated in the previous time step, we expect a higher probability that the next filament produced will also be labelled. The transitions within the two-state Markov Chain, between a labelled and an unlabelled filament, are described by the conditional probabilities shown in matrix *P* (1), from Box 1. These conditional probabilities depend on the number of labelled (*L*) and unlabelled (*U*) stem cells in the niche, on the total number *n* = *L* + *U* of stem cells, and on the probability of division *p* (see STAR Methods for derivation). The model assumptions are summarised in Box 1.

**Figure 3:**
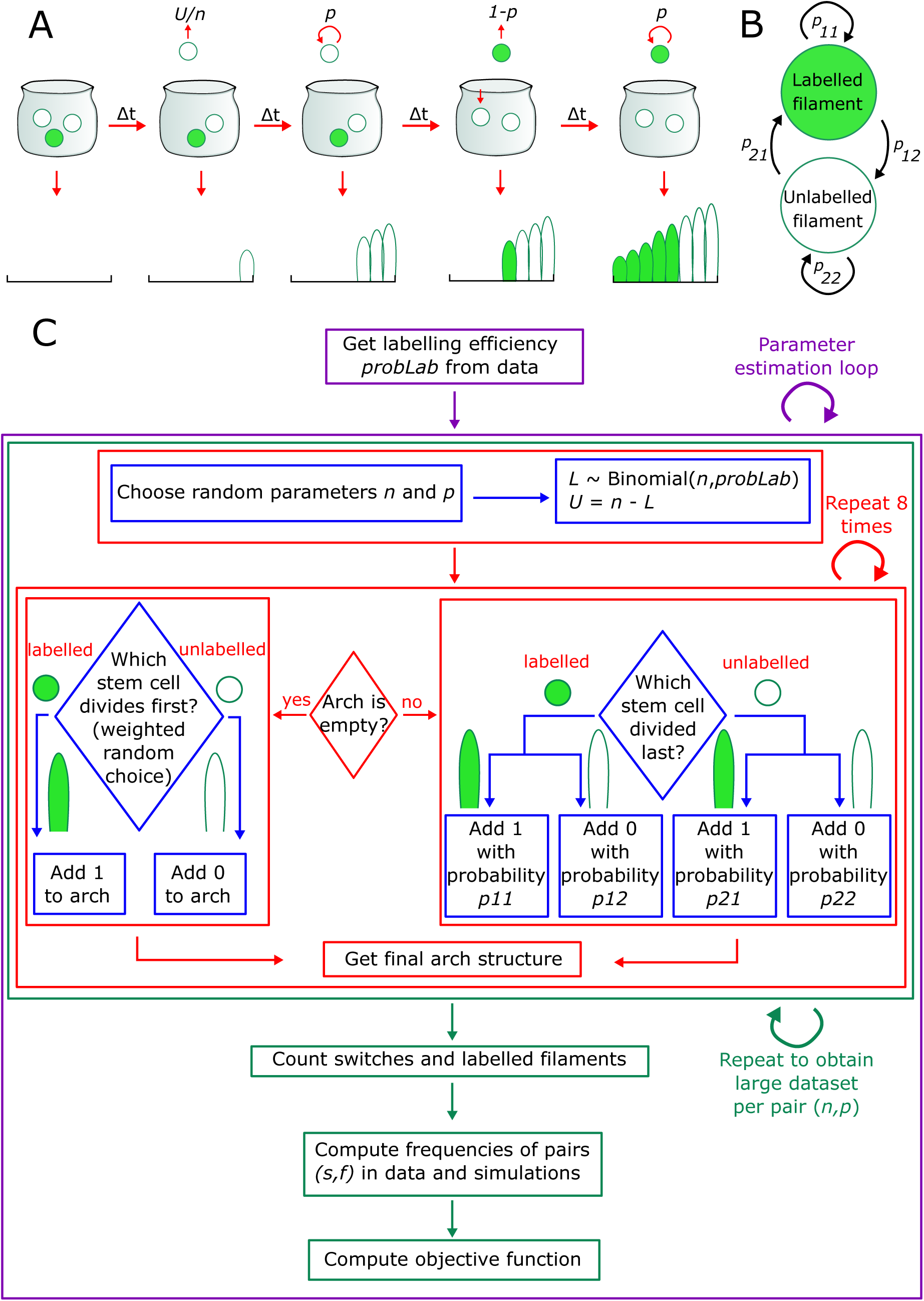
Approach for the heterogeneous model: **A**. Schematic of the model, according to the assumptions in Box 1. In the initial time step, a stem cell is randomly chosen for division. The probability of this cell to divide again is *p*, and if *p* > 0.5 more consecutive divisions of the same cell take place. At a later point, a new cell may be selected, with a probability 1 − *p*, and the probability of this cell to divide again is changed to *p*. This process continues until 8 filaments are generated. **B**. Two-state Markov Chain diagram, describing the transitions between filaments of labels 1 and 0 within the mini-arch, via the probabilities *p*_11_, …, *p*_22_. These probabilities are computed in (1). **C**. Flowchart of the stochastic algorithm presented in the main text and explained in Box 2.

#### Box 1.

**Model assumptions**

The model presented below is based on the following assumptions:

◊ One labelled stem cell (1) in the peripheral niche produces a labelled filament, while one unlabelled stem cell (0) gives rise to an unlabelled filament.
◊ The number of labelled stem cells in the niche, *L*, depends on the labelling efficiency *probLab* and on the total number of stem cells in the niche *n*: *L* ∼ Binomial(*n, probLab*), and the number of unlabelled stem cells is *U* = *n* − *L*.
◊ The cell which has just divided will divide again in the next time step with probability *p*, where the time steps correspond to division events and have random durations.

The probabilities for the transitions from one filament to another, with varying labelling status, for *n* ≥ 2 and *p* ∈ [0.5, 1), are found in the matrix *P*, with entries *p*_*ij*_ corresponding to having a filament of label *j* following a filament of label *i* in the mini-arch, with *i, j* ∈ {1, 2}, 1-labelled and 2-unlabelled.

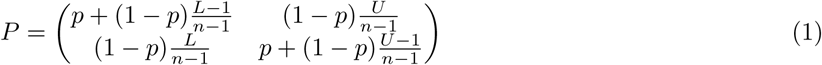

**Table B1:** Summary of parameter notations

| Parameter | Domain | Description |
| --- | --- | --- |
| $L$ | $\mathbb{N}$ | Number of *labelled* stem cells in the niche |
| $U$ | $\mathbb{N}$ | Number of *unlabelled* stem cells in the niche |
| $n = L + U$ | $\mathbb{N}$ | Total number of stem cells in the niche |
| $probLab$ | $(0, 1)$ | Labelling efficiency |
| $p$ | $[0.5, 1)$ | Probability of division |
| $s$ | $\{0, 1, \dots, 7\}$ | Number of switches within a 8-filament long mini-arch |
| $f$ | $\{0, 1, \dots, 8\}$ | Number of labelled filaments within a mini-arch |

To simulate the previously deduced heterogeneity hypothesis, a stochastic algorithm was developed, which generates a data set of *in silico* mini-arches, to compare to those from the experimental data. The two are compared by inspecting the frequencies of observing each of the pairs (*s, f*) of switches and labelled filaments. In accordance with the assumptions presented in Box 1 and based on the conditional probabilities in (1), the algorithm shown in the flowchart from Fig. 3C and outlined in Box 2 was implemented.

#### Box 2.

**A stochastic algorithm for heterogeneous stem cell division**

1. The algorithm starts by approximating the labelling efficiency based on the data, as explained by Stolper et al. (2019) and in the STAR Methods.
2. With this value at hand, the main part consisting of the stochastic simulations begins, which is visualized through the big green rectangle in the flowchart.
  a. For each mini-arch to be simulated, the program chooses random parameters *n* and *p* based on which the mini-arch will be filled with filaments, i.e. the 8-cell-long array will be filled with binary values.
  b. Out of the total number *n* of stem cells of a particular fate, the number of labelled ones, *L* ∼ Binomial(*n, probLab*) and the remaining unlabelled cells *U* = *n* − *L*. Then the *in silico* mini-arch generation begins.
    i. One starts with an empty array representing the mini-arch before any filament has been generated.
    ii. The first stem cell to divide and generate a filament is selected by a weighted random choice of whether to add a 1 or a 0 to the empty array, with weights given by probabilities of choosing a labelled cell 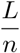, or an unlabelled cell 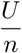, respectively.
    iii. After the first entry in the array has been added, the following ones depend on the previously inserted value. The previous dividing stem cell will divide again with probability *p*, so the labelling status of the new dividing stem cell depends on that of the previous dividing cell via the conditional probabilities (1).
    iv. The procedure stops when the array is filled with 8 entries - one virtual mini-arch.
  c. Items (i)-(iv) are repeated multiple times to generate a large table of simulated mini-arches for each pair (*n, p*) of parameters.
3. Once the simulated dataset is obtained, the pair (*s, f*) is computed for each virtual mini-arch, as it was previously done for the experimental data.
4. Subsequently, the frequency of observing the pair (*s, f*) is calculated.
5. These frequencies together with those from experimental data are used to construct an objective function

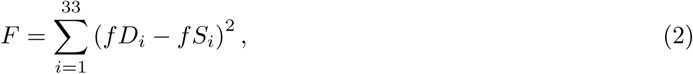

where *f D*_*i*_ and *f S*_*i*_ are the frequencies in the data and the simulation results of observing pair *i* ∈ Π, with Π = {(0, 0), (0, 8), …, (7, 4)} the set of the 33 possible pairs (*s, f*) (Supplementary Table S4). The objective function is used within the parameter estimation, to find the best parameters *n* and *p*.

The algorithm fills an initially empty array with eight binary entries, corresponding to the filaments in a miniarch, by taking into account the previously added value. A large data set of simulated mini-arches is produced. The algorithm is repeated inside a parameter estimation loop, so it is run multiple times with different starting guesses for an array of values of parameters *n* ∈ {1, …, 10}, *p* ∈ [0.5, 1) for which the objective function (2) is computed to be minimised. The objective function is a sum of square differences between the frequencies in the experimental and simulated data, of observing a pair (*s, f*) ∈ Π, with Π = {(0, 0), (0, 8), …, (7, 4)} the set of 33 possible pairs (see STAR Methods and Supplementary Table S4 for details). Finally, the best parameters *n* and *p* are obtained, which provide the most accurate fit between the simulation results and the data, performed by minimising the objective function. For this purpose, the *Mathematica* routine NMinimize was used, employing the Nelder-Mead method.

Furthermore, an analytical approach can be employed, to skip the step of simulated data generation, such that instead of computing the approximate frequencies of observing a pair (*s, f*) in the respective data, exact probabilities of each such event can be directly calculated. By using the entries of the probability transition matrix (1) together with the initial distribution 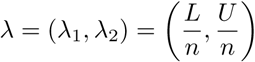, the analytical probabilities of observing a certain pair (*s, f*) of switches and labelled filaments in the model, for each parameter pair (*n, p*), are computed as a sum of probabilities of possible Markov Chain trajectories producing the required number of switches and labelled filaments. For example, the probabilities of producing an entirely unlabelled mini-arch (with 0 switches and 0 labelled filaments) and an entirely alternating mini-arch (with 7 switches and 4 labelled filaments) read:

◊ 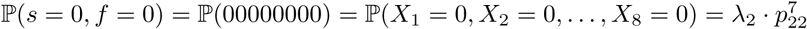 and
◊ 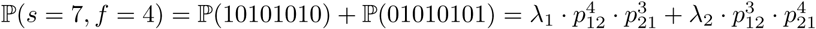.

Similarly, this method can be employed for each of the 33 possible pairs (*s, f*) to obtain a list of probabilities with entries corresponding to each pair, depending on the number of stem cells in the niche *n*, the probability of division *p* and the labelling efficiency *probLab*. As in the previous numerical method, a sum-of-squares objective function is minimized in order to obtain the best parameter set. In the analytical approach, the frequencies *f S*_*i*_ in (2) (Box 2) are replaced by the exact probabilities of observing a pair (*s, f*) ∈ Π, which can be computed as summarized above.

In order to obtain an initial general overview on the parameters’ influence on the data, the logarithm of the objective function is plotted against the two parameters, *n* and *p* (Fig. 4A). These plots suggest that few stem cells are sufficient to generate filaments in the branchial arches, as long as their probability of division is high, indicating a highly heterogeneous division behaviour that corresponds to activation and quiescence phases. In the case of pattern 4, there is a higher variability in the values for both *n* and *p*, which we attribute to practical non-identifiability - see Discussion. In pattern 2, the probability of division *p* seems to be smaller than in the other patterns, suggesting that the activation phases of SCs of fate 2 are shorter. Subsequently, the parameters *n* and *p* are estimated by minimising the respective objective functions, as discussed before.

**Figure 4:**
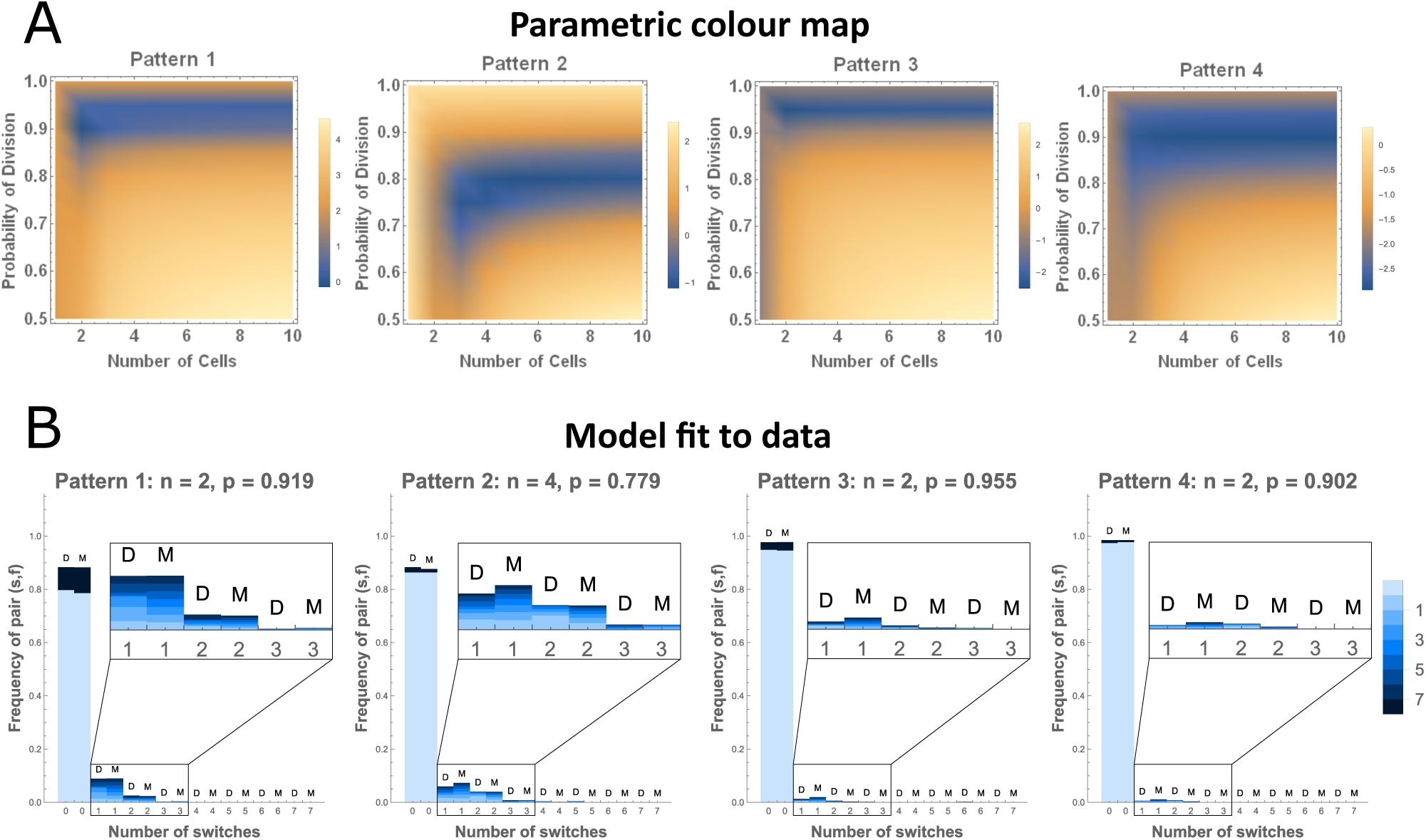
Results of the heterogeneous model: **A**. Parametric colour maps representing the logarithms of the objective functions (scaled by 103) with respect to the number of cells *n* and probability of division *p*. The smaller the objective function, the better the fit to the data. Hence, the coordinates in the (*n, p*) plane of the darkest point represent the best parameter values. Plots correspond to patterns 1-4, from left to right. **B**. Comparisons between frequencies and probabilities of observing a pair (*s, f*) in experimental data (D) and model results (M), respectively, for each of the four patterns (1-4 from left to right). The *x*-axis represents the number of switches and the colour code corresponds to the number of labelled filaments (from light blue - zero to dark blue - eight). Parameters were estimated by minimizing the objective function computed according to the analytical probabilities, and can be read in the titles of each plot.

Both methods provide agreeing results, with similar parameter values, as expected. Fig. 4B shows plots for each pattern, using the parameter values estimated with the analytical approach. These plots, based on estimated parameters, show a good fit between the model and data and suggest that only few stem cells for each pattern are participating in organ growth (2-4 stem cells, depending on the pattern), with high probabilities of division (≈ 0.9 for most). Some differences can be observed, for example, in the ratio between fully labeled and fully unlabelled mini-arches (dark and light blue in the zero switches bar), which is influenced by the labelling efficiency, as previously mentioned. This minor mismatch comes from the approximation of the labelling efficiency from the data. Even though the method is based on realistic assumptions, in some cases it does not provide a close enough value to the true one, because the dataset is not sufficiently large for this task (*N* = 470 mini-arches). If a larger data set were available (*N* = 104 mini-arches), a much better approximation could be obtained. Nevertheless, the bars for data (D) and model (M) have equal heights in most cases and provide a good fit, indicating that few but highly heterogeneous stem cells are enough to contribute filaments to the medaka gills during the entire post-embryonic and adult life.

## Discussion

This study has presented two alternative methods for determining the numbers and functional heterogeneities among stem cells during post-embryonic gill growth: a numerical approach *via* stochastic simulations, which generates a large data set of simulated mini-arches for comparison to experimental data, and a more abstract analytical approach, which skips the step of *in silico* data generation, is exact and less time-consuming. The two methods are based on the same assumptions and provide equivalent results. These methods contributed to the discovery of novel insights into the behaviour of the stem cells responsible for building, growing and maintaining the respiratory organ of fish: (i) few branchial arch stem cells are participating in the post-embryonic and adult organ growth and, more importantly, (ii) these stem cells are functionally heterogeneous in their division behaviour, in the sense that they follow phases of activation and quiescence, such that once a stem cell has divided to generate a filament, it becomes active and divides multiple times thus creating more clonal filaments, before becoming quiescent and allowing another stem cell of the same type to take over the task of filament generation.

This activation/quiescence behaviour is valid for all four fate-restricted stem cells, corresponding to the four possible patterns observed in filaments. In all cases, therefore, the heterogeneity parameter *p* describing the probability of division is very high. Stem cells of pattern 2 has a smaller probability of division, relative to the other, which suggest that their activation phases are shorter, and this comes together with a higher number of SCs of the second type, which participate in organ growth. In pattern 4 more variability is observed (Fig. 4A), which is caused by the low labelling efficiency in the case of the fourth stem cell type. This low labelling efficiency may also be a consequence of the fact that Pattern 4 is difficult to spot when mixed patterns are present, being hidden beneath other more prominent patterns (e.g. Pattern 2). A suitable method to overcome this issue would be to label the different types of cells (patterns) with different colours, but such method is currently not available with the required cellular-resolution for the gill system.

The results of this study improve our understanding of the coordination between the filament-generating stem cells, which get recruited as an ensemble to a newly forming filament. The aspect of how this coordination is accomplished remains an open question, but our studies have shown that approximately equal numbers of the fate-restricted stem cells are responsible for organ growth, which supports the mechanism of maintaining the ratio of cell types within a filament, guaranteeing its proper development. The alternating activation and quiescence phases, have also been recently suggested in other systems, for example in neurogenesis (Harris et al., 2020; Urbán et al., 2016; Ziebell et al., 2018; Basak et al., 2018), where it is speculated that proliferating stem cells return to quiescence into a pool of temporary quiescent cells, which is separate from the main dormant stem cells. This behaviour can be thought of as a defence mechanism against the possible steps which can fail during division, considering the small number of stem cells which carry the responsibility of filament generation and thus, organ growth. We do not exclude that in addition to the respective 2-4 stem cells driving the organ growth, more stem cells of each particular fate reside in the niche, but according to our model and experimental data the majority of them should be dormant, only becoming active when one of the main stem cells fails and dies.

## Supporting information

Key Resource Table

## Acknowledgements

This project is supported by the German Research Foundation (DFG), as part of the Collaborative Research Centre SFB873 “Maintenance and Differentiation of Stem Cells in Development and Disease”. The project is a collaboration between groups B09 “Mathematical Modeling of Stem Cell Renewal and Differentiation” led by AM-C and A11 “Mechanisms of Niche – Stem Cell Unit Origin and Maintenance” led by LC.

## Author Contribution

D-PD: Model development, Algorithm implementation, Formal analysis, Data curation, Interpretation of the modelling results, Manuscript preparation - original draft;

JS: Data acquisition and curation, Manuscript preparation - review and editing;

LC: Conceptualization, Supervision, Data acquisition and curation, Manuscript preparation - review and editing;

AM-C: Conceptualization, Supervision, Model development, Formal analysis, Interpretation of the modeling results, Manuscript preparation - review and editing

## STAR Methods

### EXPERIMENTAL MODEL AND SUBJECT DETAILS

#### Animal husbandry and ethics

All fish stocks of *Oryzias latipes* (medaka) were maintained according to the local animal welfare laws (Tierschutzgesetz §11, Abs. 1, Nr.1) and the European Union animal welfare guidelines. Fish were maintained and raised in a constant recirculation system at 28°C cycling between 14 hours of light and 10 hours of darkness (Tierschutzgesetz §11, Abs. 1, Nr.1, Haltungserlaubnis AZ35-9185.64 and AZ35-9185.64/BH KIT). Fish lines being used in this study include wild-type medaka (Cab, medaka Southern population strain) and transgenic fish of the Gaudí living toolkit (Centanin et al., 2014): Gaudí^*Ubiq*.*iCre*^, expressing a tamoxifen inducible Cre-recombinase, Gaudí^*HSP*70.*A*^, expressing a heat-shock inducible CRE-recombinase and Gaudí^*RSG*^, containing a genetic cassette that switches from a ubiquitous expression of a red fluorescent protein (RFP) to a nuclear green signal (nGFP) upon recombination.

### METHOD DETAILS

#### Generation of clones

Clonal data was generated via lineage tracing analysis. Genetic recombination in double transgenic fish (Gaudí^*HSP*70.*A*^, Gaudí^*RSG*^) was induced via heat-shock. Embryos were staged according to Iwamatsu (2004) and heat-shocked at stages 20, 24, 29 32, 34 or 37 using embryo rearing medium (ERM) at 42°C and transferred to 37°C for 1 to 3 hours and raised until adulthood.

Genetic recombination was induced via tamoxifen treatment in Gaudí^*Ubiq*.*iCre*^ Gaudí^*RSG*^ double transgenic fish at stage 36. Embryos were kept in ERM containing tamoxifen (T5648 Sigma, 5*µ*M final concentration) for 3 hours, rinsed multiple times with fresh ERM to ensure removal of residual tamoxifen, and placed in a tank until they reach adulthood. Fish that resulted in a high recombination efficiency (i.e. entire branchial arch labelled) were not used in the analysis.

#### Staining protocol and imaging

All fish were euthanised using a 2mg/ml Tricaine solution (Sigma-Aldrich, A5040-25G), fixed in 4% PFA/PTW at 4°C overnight and the entire gills were micro-dissected to continue with the staining protocol. To permeabilise the tissue, gills were kept in acetone at −2°C for 10 minutes. After blocking with goat serum for 1 hour at room temperature, GFP staining (Rabbit a-GFP, Invitrogen, 1:750) was performed overnight at 4°C. The secondary antibody (Alexa 488 a-Rabbit, Invitrogen, 1:500) was incubated together with DAPI (final concentration: 5*µ*g/*µ*l) for 2 hours at room temperature. Gills were separated into single branchial arches and mounted in glycerol 50% between cover slides (Stolper et al., 2019). Whole gills were imaged using an Olympus MVX10 microscope connected to a Leica DFC500. On confocal resolution, branchial arches were imaged using Leica TCS SP8 and SP5 II microscopes. Image analysis and stitching was performed in Fiji.

### QUANTIFICATION AND STATISTICAL ANALYSIS

#### Experimental data adjustment

The experimental data consisted of a table with rows of unequal lengths, each of which represented one branchial arch. Each such array contained values describing the labelling status and respective pattern or combination of patterns, for each filament in the arch. An unlabelled filament was recorded as an element *i* = 0, a filament presenting one pattern was denoted by *i* ∈ {1, 2, 3, 4}, while a filament presenting mixed patterns was presented as a multiple digit number, with digits recording the patterns present, e.g. for a combination of two patterns within one filament the options are 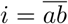, with *a, b* ∈ {1, 2, 3, 4}, *a < b* (Table S1, excerpt from experimental data). A total of *N* = 340 branchial arches were quantified. From these, only the branchial arches/arrays of length ≥ 25 were considered, resulting in a total of *N* = 235 rows. Subsequently, four separate data sets were created, one for each pattern, by only considering the filaments presenting that specific pattern as labelled (1), and the other cases as unlabelled (e.g. in the adjusted data for Pattern 1, the filaments with patterns 2, 3 and 4, together with the unlabelled filaments are recorded as 0) - see Table S2, data excerpt adjusted for each pattern. For all patterns, from each branchial arch, the 8 peripheral filaments from each side were selected, i.e. the first 8 and last 8 entries of each array. All such mini-arches were arranged so that they start with the oldest filament, i.e. the mini-arrays containing the first 8 entries from the original array were flipped. The final four data sets used for the model, one for each pattern, each consisted of a 8 × 2*N* = 8 × 470 matrix (Table S3, final adjusted excerpt).

#### Labelling efficiency estimation

The labelling efficiency was estimated for the entire experimental data set by employing a combinatorial approach. The labelling efficiency would indicate how many stem cells are labelled out of a large pool of SCs. To keep in mind is the fact that a labelled filament is produced by a labelled stem cell, so if one looks at the oldest post-embryonic filament, one finds the label of the first SC which divided in that specific branchial arch (on each side). We look at the oldest filament because, in this way, we avoid the influence of the heterogeneity parameter. Accordingly, by counting the number of labelled first filaments (i.e. oldest) out of the total number of branchial arches, we obtain an approximation of the average labelling efficiency across the entire data set. We thus obtain the following formula, where *nL* is the number of mini-arches which start with a labelled filament, i.e. the number of arrays that start with a non-zero (1) entry; and *N* is the total number of rows in the adjusted data set.

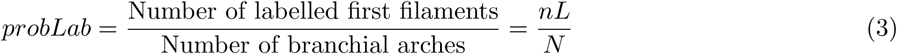

Accordingly, for each pattern the labelling efficiency *probLab* reads as in Table 1.

**Table 1:** Labelling efficiency for each pattern.

| Pattern | Pattern 1 | Pattern 2 | Pattern 3 | Pattern 4 |
| --- | --- | --- | --- | --- |
| $probLab$ | 0.1553 | 0.0638 | 0.0426 | 0.0149 |

#### Transition probabilities computation

First, note that we choose *p* ∈ [0.5, 1) based on our previous heterogeneity hypothesis. A probability of division of *p* = 0.5 corresponds to an entirely functionally homogeneous system, in which the probability of the previously diving cell to divide again is equal to that of another random cell of the same type to take over. A probability of division *p* > 0.5 corresponds to our hypothesis of a functionally heterogeneous system with activation and quiescence phases. In addition, *p <* 1, since *p* = 1 would be equivalent to a case where the entire mini-arch is created by one stem cell, case which we proved infeasible (Fig. 2D and Stolper et al. (2019)).

For computing the transition probabilities of the Markov process, recall the formula for conditional probabilities

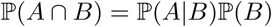

Suppose *i* is the cell which has just divided and *j* the cell about to divide. Further, denote by *cL* the event of choosing a labelled cell, and by *cU* the event of selecting an unlabelled cell. A transition *L* → *L* i.e. a labelled filament followed by another labelled one, corresponds to a case where either the previously diving cell *i* was labelled and it divides again (*j* = *i*), or if another labelled cell is selected (*j* ≠ *i*, with *i, j* labelled). All transition probabilities can be similarly considered and recorded in the transition probability matrix *P* with entries (4) for *n* ≥ 2, recalling that *p* = ℙ(*j* = *i*) irrespective of the labelling status. For *n* = 1, the matrix *P* = *I*_2_ the identity matrix, but this case is not expected (Fig. 2D and Stolper et al. (2019)).

**Entries of the transition probability matrix *P*, for *n* ≥ 2:**

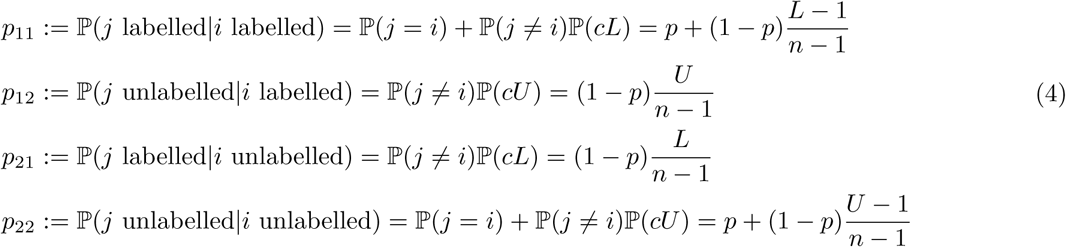

#### Objective function definition

The objective function describes a measure of the “distance” between the model and the experimental data. In this discrete system, the data are quantified based on the (*s, f*) pairs, so the objective function compares the frequencies of observing such a pair, in the data and in the model. There exist 33 possible pairs describing a mini-arch since, out of the total 8 × 9 pair combinations, most are infeasible due to configuration dependency constraints. For example, there exists no mini-arch corresponding to a pair (*s, f*) = (1, 8) because in an entirely labelled mini-arch (*f* = 8) no switches exits (*s* = 0). The set Π of all possible pairs was obtained by implementing a short piece of code investigating each mini-arch configuration (256 configurations) - see Table S4 for all possible configurations and details. The objective function is thus constructed as a sum of square differences

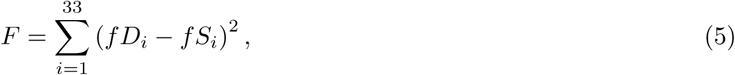

where *fD*_*i*_ and *fS*_*i*_ are the frequencies in the data and the simulation results of observing pair *i* ∈ Π, with Π = {(0, 0), (0, 8), …, (7, 4)} the set of the 33 possible pairs (*s, f*).

## DATA AND CODE AVAILABILITY

The experimental data and the code generated during this study are available on GitHub: https://github.com/dpdanciu/Gill_SCnumbers-heterogeneities.

## KEY RESOURCES TABLE

