## Supplementary material for "Identifying stem cell numbers and functional heterogeneities during post-embryonic organ growth": Key Resource Table

**KEY RESOURCES TABLE**

| REAGENT or RESOURCE | SOURCE | IDENTIFIER |
| --- | --- | --- |
| Antibodies | | |
| a-EGFP (Rabbit IgG polyclonal) | Invitrogen (Thermo Fischer) | CAB4211; RRID: [AB_10709851](https://scicrunch.org/resolver/AB_10709851) |
| Alexa 488 Goat a-Rabbit | Invitrogen (Thermo Fischer) | A-11034 |
| Chemicals, Peptides, and Recombinant Proteins | | |
| tamoxifen | Sigma-Aldrich | T5648 |
| tricaine | Sigma-Aldrich | A5040-25G |
| DAPI | Roth |  |
| Experimental Models: Organisms/Strains | | |
| Wild type *Oryzias latipes,* Cab |  |  |
| Transgenic *Oryzias latipes,* GaudíUbiq.iCre | Centanin et al., 2014 |  |
| Transgenic *Oryzias latipes,* GaudíHsp70.A | Centanin et al., 2014 |  |
| Transgenic *Oryzias latipes,* GaudíRSG | Centanin et al., 2014 |  |
| Software and Algorithms | | |
| Fiji | <https://fiji.sc/> |  |
| Mathematica 12.0 | Wolfram Research |  |
| Other | | |
