## Supplementary Table 4 for "Identifying stem cell numbers and functional heterogeneities during post-embryonic organ growth"

**Table S4**: By inspecting the total number of binary sequences within an 8-entries long array (i.e. the 256 binary numbers of up to 8 digits, those with less digits being filled with 0s at the beginning), and determining the number of switches and labelled filaments for each, we obtain 33 possibilities for the pairs (s,f), with s switches and f labelled filaments. The possible configurations for each such pair are presented in the table below. The code for computing all pairs and arch configurations was implemented in Mathematica.

| **Pair** | ***s*** | ***f*** | **Mini-arch configurations** |
| --- | --- | --- | --- |
| **1** | **0** | **0** | 00000000 |
| **2** |  | **8** | 11111111 |
| **3** | **1** | **1** | 10000000, 00000001 |
| **4** |  | **2** | 11000000, 00000011 |
| **5** |  | **3** | 11100000, 00000111 |
| **6** |  | **4** | 11110000, 00001111 |
| **7** |  | **5** | 11111000, 00011111 |
| **8** |  | **6** | 11111100, 00111111 |
| **9** |  | **7** | 11111110, 01111111 |
| **10** | **2** | **1** | 01000000, 00100000, 00010000, 00001000, 00000100, 00000010 |
| **11** |  | **2** | 10000001, 01100000, 00110000, 00011000, 00001100, 00000110 |
| **12** |  | **3** | 11000001, 10000011, 01110000, 00111000, 00011100, 00001110 |
| **13** |  | **4** | 11100001, 11000011, 10000111, 01111000, 00111100, 00011110 |
| **14** |  | **5** | 11110001, 11100011, 11000111, 10001111, 01111100, 00111110 |
| **15** |  | **6** | 11111001, 11110011, 11100111, 11001111, 10011111, 01111110 |
| **16** |  | **7** | 11111101, 11111011, 11110111, 11101111, 11011111, 10111111 |
| **17** | **3** | **2** | 10100000, 10010000, 10001000, 10000100, 10000010, 01000001, 00100001, 00010001, 00001001, 00000101 |
| **18** |  | **3** | 11010000, 11001000, 11000100, 11000010, 10110000, 10011000, 10001100, 10000110, 01100001, 01000011, 00110001, 00100011, 00011001, 00010011, 00001101, 00001011 |
| **19** |  | **4** | 11101000, 11100100, 11100010, 11011000, 11001100, 11000110, 10111000, 10011100, 10001110, 01110001, 01100011, 01000111, 00111001, 00110011, 00100111, 00011101, 00011011, 00010111 |
| **20** |  | **5** | 11110100, 11110010, 11101100, 11100110, 11011100, 11001110, 10111100, 10011110, 01111001, 01110011, 01100111, 01001111, 00111101, 00111011, 00110111, 00101111 |
| **21** |  | **6** | 11111010, 11110110, 11101110, 11011110, 10111110, 01111101, 01111011, 01110111, 01101111, 01011111 |
| **22** | **4** | **2** | 01010000, 01001000, 01000100, 01000010, 00101000, 00100100, 00100010, 00010100, 00010010, 00001010 |
| **23** |  | **3** | 10100001, 10010001, 10001001, 10000101, 01101000, 01100100, 01100010, 01011000, 01001100, 01000110, 00110100, 00110010, 00101100, 00100110, 00011010, 00010110 |
| **24** |  | **4** | 11010001, 11001001, 11000101, 10110001, 10100011, 10011001, 10010011, 10001101, 10001011, 01110100, 01110010, 01101100, 01100110, 01011100, 01001110, 00111010, 00110110, 00101110 |
| **25** |  | **5** | 11101001, 11100101, 11011001, 11010011, 11001101, 11001011, 10111001, 10110011, 10100111, 10011101, 10011011, 10010111, 01111010, 01110110, 01101110, 01011110 |
| **26** |  | **6** | 11110101, 11101101, 11101011, 11011101, 11011011, 11010111, 10111101, 10111011, 10110111, 10101111 |
| **27** | **5** | **3** | 10101000, 10100100, 10100010, 10010100, 10010010, 10001010, 01010001, 01001001, 01000101, 00101001, 00100101, 00010101 |
| **28** |  | **4** | 11010100, 11010010, 11001010, 10110100, 10110010, 10101100, 10100110, 10011010, 10010110, 01101001, 01100101, 01011001, 01010011, 01001101, 01001011, 00110101, 00101101, 00101011 |
| **29** |  | **5** | 11101010, 11011010, 11010110, 10111010, 10110110, 10101110, 01110101, 01101101, 01101011, 01011101, 01011011, 01010111 |
| **30** | **6** | **3** | 01010100, 01010010, 01001010, 00101010 |
| **31** |  | **4** | 10101001, 10100101, 10010101, 01101010, 01011010, 01010110 |
| **32** |  | **5** | 11010101, 10110101, 10101101, 10101011 |
| **33** | **7** | **4** | 10101010, 01010101 |
